# Modulation of Rapid Visual Responses during Reaching by Multimodal Stimuli

**DOI:** 10.1101/570796

**Authors:** Isabel S Glover, Stuart N Baker

**Author notes:** **Disclosures:** No conflicts of interest, financial or otherwise, are declared by the authors. **Author Contributions:** I.S.G. and S.N.B. conceived and designed research; I.S.G. performed experiments and analyzed data; I.S.G. and S.N.B. interpreted results of experiments; I.S.G. prepared figures and drafted manuscript; I.S.G. and S.N.B. edited and revised manuscript; I.S.G. and S.N.B. approved final version of manuscript. **Funding**: This study was funded by The Reece Foundation.

## Abstract

The reticulospinal tract plays an important role in primate upper limb function, but methods for assessing its activity are limited. One promising approach is to measure rapid visual responses (RVRs) in arm muscle activity during a visually-cued reaching task; these may arise from a tecto-reticulospinal pathway. We investigated whether changes in reticulospinal excitability can be assessed non-invasively using RVRs, by pairing the visual stimuli of the reaching task with electrical stimulation of the median nerve, galvanic vestibular stimulation or loud sounds, all of which are known to activate the reticular formation.

Surface electromyogram recordings were made from the right deltoid of healthy human subjects as they performed fast reaching movements towards visual targets. Stimuli were delivered up to 200ms before target appearance and RVR was quantified as the EMG amplitude in a window 75-125ms after visual target onset. Median nerve, vestibular and auditory stimuli all consistently facilitated the RVRs, as well as reducing the latency of responses. We propose that this reflects modulation of tecto-reticulospinal excitability, suggesting that the amplitude of RVRs can be used to assess changes in brainstem excitability non-invasively in humans.

**New & Noteworthy:** Short latency responses in arm muscles evoked during a visually-driven reaching task have previously been proposed to be tecto-reticulospinal in origin. We demonstrate that these responses can be facilitated by pairing the appearance of a visual target with stimuli that activate the reticular formation – median nerve, vestibular and auditory stimuli. We propose that this reflects non-invasive measurement and modulation of reticulospinal excitability.

## Introduction

The reticulospinal tract (RST) projects to motoneurons innervating both distal and proximal muscles in primates (Davidson and Buford 2006; Davidson and Buford 2004; Riddle et al. 2009) and increasing evidence supports its role in upper limb function (Baker 2011), from gross reaching (Schepens and Drew 2006; 2004) to precise finger movements (Carlsen et al. 2009; Honeycutt et al. 2013; Soteropoulos et al. 2012). Given the potential of this pathway to mediate functional recovery (Baker 2011; Baker et al. 2015), it would be highly desirable to develop means of assessing and modulating RST activity in humans.

For the corticospinal tract, considerable progress has been made by using transcranial magnetic stimulation (TMS) to excite motor-evoked potentials (MEPs) in contralateral muscles. By conditioning TMS with a prior stimulus and measuring whether the MEP is facilitated, it is possible to determine whether that stimulus can influence corticospinal excitability (e.g. Furubayashi et al. 2000; Tokimura et al. 2000). If we are to use a similar approach for the RST, it is first necessary to find a way of generating a test response which is likely to be mediated mainly by the RST. MEPs in muscles ipsilateral to the stimulus (Ziemann et al. 1999), and the long-latency stretch reflex in proximal muscles (Foysal et al. 2016) may both have potential in this regard. A further possibility exploits the projections from the deep layers of the superior colliculus to the reticular formation (RF; Grantyn and Grantyn 1982; Illert et al. 1978).

Several lines of evidence support a role for this tecto-reticulospinal pathway in upper limb movement. Cells within the superior colliculus and the underlying RF modulate their discharge with arm movements (Stuphorn et al. 1999; Werner 1993), and microstimulation of both areas can evoke activity in proximal arm muscles (Philipp and Hoffmann 2014). Furthermore, lesion studies in cats have identified the tecto-reticulospinal tract as an important substrate in mediating early responses to visual perturbations (Alstermark et al. 1987). In humans, fast reaching movements made towards visual targets evoke short-latency EMG responses in proximal muscles (Pruszynski et al. 2010). These rapid visual responses (RVRs) are stimulus-locked (Pruszynski et al. 2010) and relatively independent of volition (Gu et al. 2016). It has therefore been suggested that they may bypass the cortex, and are mediated by the tecto-reticulospinal tract. If this is the case, it suggests that measurement of RVRs could provide an assessment of the excitability of the RST, in the same way that MEPs allow insight into corticospinal function.

In addition to visual information from the superior colliculus, the RF also receives sensory information from peripheral afferents (Leiras et al. 2010), auditory stimuli (Irvine and Jackson 1983) and the vestibular system (Ladpli and Brodal 1968; Peterson and Abzug 1975). This extensive convergence of multisensory information should provide ample opportunities to modulate RST excitability. In this study, we therefore assessed whether pairing visual target appearance with stimuli known to activate the RF could modulate the RVRs generated during a reaching task. We were able to demonstrate RVR facilitation by stimulation of peripheral afferents, the vestibular system, and loud sounds, in a manner consistent with convergence within the brainstem.

## Methods

### Subjects

Eight subjects participated in each of three separate experiments, which tested the effect of different conditioning stimuli: electrical stimulation of the median nerve (age: 19.9 ± 1.7 years; 1 female), galvanic stimulation of the vestibular system (age: 19.9 ± 1.9 years; 1 female), and a loud auditory stimulus (age: 22.7 ± 3.7 years; 4 female). Some subjects participated in multiple experiments. All subjects were right-handed, had no history of neurological disorders, and provided written informed consent to participate in the study. All procedures were approved by the local ethics committee and the study complied with the Declaration of Helsinki.

### EMG Recordings

Surface EMG recordings were made from the right lateral deltoid and pectoralis major (PM). Two silver/silver chloride electrodes (Kendall H59P, Medcat) were placed on the skin overlying each muscle along the direction of the muscle fibers. In the median nerve and vestibular protocols, intramuscular EMG was also recorded using custom-made fine-wire electrodes (7 stranded stainless steel wire coated in Teflon insulation; Advent Research Materials catalogue number FE6320). All EMG signals were amplified (200-10,000 gain), filtered (30Hz to 2kHz bandpass) and digitized (5kHz) for off-line analysis (CED 1401 with Spike2 software, Cambridge Electronic Design).

### Experimental Sessions

Our experimental task was based upon that reported by Pruszynski et al. (2010). Subjects grasped an ergonomically-shaped handle at the end of a manipulandum comprising two metal shafts connected to each other and a firm base by vertical revolving joints (Figure 1***A***). This permitted free movement in the horizontal plane; optical encoders on the joints allowed measurement of end point position. Subjects were comfortably seated in front of this device, and held the handle in their right hand with the elbow flexed around 90°. A video monitor and half-silvered mirror allowed the projection of targets into a plane aligned to the top of the handle. A red LED placed on the handle in this plane indicated hand position at appropriate times during each trial. Experiments were performed in the dark; the half-silvered mirror prevented subjects from seeing their own hand, so that the LED (when lit) was the only visual information available about hand position.

**Figure 1.**
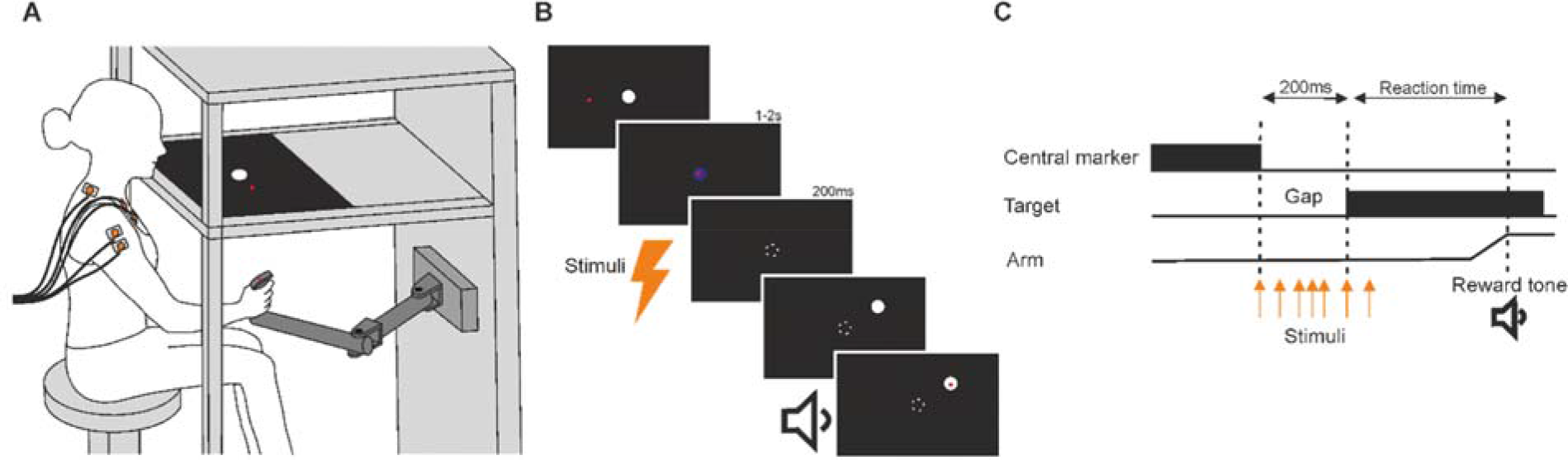
Experimental paradigm. ***A.*** Subjects made reaching movements in a horizontal plane by moving a manipulandum with their right hand. Targets were displayed on a screen and projected onto the plane of movement using a half silvered mirror that occluded view of the hand. A red LED on the handle of the manipulandum indicated position when illuminated. ***B,C.*** Each trial began with the presentation of a central marker (white circle, 1 cm radius). Subjects were required to align their hand with this; the central marker turned blue when the hand was correctly aligned. This position was maintained for a randomized period of 1-2 s. The central marker then disappeared for a fixed gap period of 200 ms and the red LED was turned off. Following the gap period, a peripheral target (white circle, 1 cm radius) appeared in one of four directions (45°, 135°, 225° or 315° relative to the right horizontal axis, 10cm from the central marker). Subjects were instructed to move to this target as quickly as possible. Once reached, the red LED turned on again, the target disappeared and the central marker reappeared indicating the start of the next trial. Subjects were provided with auditory feedback of task performance. Stimuli (loud sounds, median nerve stimulation or galvanic vestibular stimulation) were delivered between 200 ms before and 50 ms after target appearance (orange arrows). No stimuli were delivered during the control condition.

The trial sequence is outlined in Figure 1***B***. The appearance of a central marker (white circle, 1 cm radius) indicated the start of each trial. Subjects moved the handle to this marker at their own pace, placing the illuminated LED within the projected circle. Successful alignment was indicated by the circle changing color from white to blue. Subjects were required to maintain this position for a randomized period of 1-2 s, after which both the circle and LED disappeared for a gap period of 200 ms, which has been shown to decrease reaction times (Fischer and Rogal 1986; Gribble et al. 2002). The imperative stimulus consisted of a peripheral target (white circle, 1 cm radius) which appeared in one of four directions (45°, 135°, 225° or 315° relative to the right horizontal axis) at a distance of 10 cm from the central position. Subjects were instructed to make fast reaching movements to this new target. The red LED was turned on again only when the target was reached; this encouraged subjects to make ballistic rather than tracking movements. Auditory feedback was provided at the end of each trial to indicate whether the target was reached in less than 500 ms.

Subjects performed blocks of 40 trials (10 in each direction), separated by rest periods of 60 s in which the mean reaction time for the preceding block was presented on the screen. For all experiments, subjects completed a total of 960 trials (24 blocks of 40 trials).

### Stimulus Conditions

A separate experiment was performed for each of the following stimuli: electrical stimulation of the median nerve at the wrist, galvanic vestibular stimulation and loud sounds. Stimuli were delivered at five different latencies relative to the visual target appearance (median nerve: −200, −100, −50, 0, 50 ms; vestibular and auditory: −150, −100, −75, −50, 0 ms; negative latencies indicate stimuli delivered prior to target appearance). Trials with stimuli were interleaved randomly with a control (unstimulated) condition. The 24 different trial types (4 target directions x 6 stimulus conditions) were tested in an order randomized across the entire experiment.

Median nerve stimulation (500 μs pulse, Digitimer DS7A isolated stimulator) was delivered through adhesive electrodes (Kendall H59P, Medcat) placed over the right median nerve at the level of the wrist (cathode proximal). Motor threshold was assessed as the minimum intensity required to produce a twitch in the thenar muscles; stimulation during the experiment was at twice motor threshold. Galvanic vestibular stimulation (4 mA, 20 ms pulse; Digitimer DS4 isolated stimulator) was delivered through adhesive electrodes (F-RG/6, Skintact) placed over the mastoid processes (cathode left). Auditory stimuli (120 dB SPL, 20 ms duration 1 kHz sinusoidal tone) were delivered through speakers positioned in front of the subject.

Each experiment lasted approximately one hour. Task parameters including handle position, stimulus condition, target direction and reaction time were stored to disc along with EMG recordings. To prevent timing errors potentially introduced by the video display, a small white square was displayed in the corner of the video screen at the same time as the target. A photodiode was fixed to this location on the screen with opaque tape; the square was therefore not visible to the subject but the photodiode generated a clear voltage change at target appearance, which was used for trial alignment in analysis.

### Data Analysis

All data analysis was performed off-line using custom software written in MATLAB. EMG recordings were high pass filtered at 30 Hz, full-wave rectified and smoothed by convolution with a Gaussian (mean parameter μ=0 ms; width parameter σ=1 ms).

Trials were excluded from subsequent analysis if the initial movement was made in the wrong direction, defined as the first 5 mm of movement not being in the appropriate 90° arc towards the target. Trials were also excluded on the basis of movement time. This was not assessed simply as the time taken to reach the target, since it was common for subjects narrowly to miss the target and then spend considerable time searching for it, a task made difficult since they could not see their hand. Instead, for trials that were made in the correct direction, we measured the time taken to reach 10 cm from the center (the target distance). This provided a measure of movement time independent of movement accuracy. Trials with movement time exceeding 500 ms were excluded.

We observed two notable effects in the EMG traces. Firstly, there was a band of short-latency activity which resembled the visual response described by Pruszynski et al. (2010). We refer to this as the rapid visual response (RVR). The amplitude of the RVR was calculated as the area under the curve above baseline EMG between 75 and 125 ms, as this window encompasses the range of values reported in the literature (Gu et al. 2018; Gu et al. 2016; Pruszynski et al. 2010). The RVR amplitude was normalized by expressing it as a percentage of the mean total EMG activity for the control (unstimulated) condition. Because stimuli could sometimes change the total EMG activity, we also calculated RVR size as a percentage of the total EMG activity measured on the same single trial. Total EMG activity for each trial was calculated as the area under the curve, above baseline EMG, measured from the target appearance until the time at which target distance was reached. Baseline EMG activity for each trial was measured in the 500 ms preceding the gap period (i.e. 700 to 200 ms before target appearance).

The second effect observed in the EMG traces was a latency shift with stimulation. Latencies were measured from averaged traces for a given condition. EMG onset latency was defined as the time point at which EMG activity exceeded a threshold value of two standard deviations above mean baseline EMG activity for at least 50 ms. Latencies are expressed relative to the target onset time, such that negative values represent an increase in EMG activity prior to the target appearance.

The effect of stimulus and target direction on RVR size and EMG latency was assessed using two-way repeated measures ANOVAs. The effect of the stimuli on total EMG activity, time to target, time to target distance and error rates was assessed using one-way repeated measures ANOVAs. Post-hoc analysis was performed with t-tests. The significance threshold was set at P<0.05.

Similar trends were observed for recordings from the deltoid and pectoralis major muscles and for surface and intramuscular EMG, although the results were clearest in the surface data from deltoid. This is possibly due to the difficulty of obtaining high quality recordings from pectoralis major in female subjects, and the broader sampling of muscle activity for surface compared to intramuscular EMG. In this paper, we therefore report only the findings using surface recordings from deltoid.

## Results

All subjects successfully completed the protocol. The procedure described in Methods to exclude inaccurate and slow trials led to a total of 84.3 ± 7.8 % of trials being included for the median nerve protocol, 84.5 ± 6.9 % for the vestibular protocol and 73.5 ± 12.7 % for the auditory protocol (mean ± SD across subjects).

### Effects of Stimuli on RVR Amplitude

Relative to the control condition, the stimuli appeared to facilitate the RVR (median nerve: Figure 2; vestibular: Figure 3; auditory: Figure 4). To quantify this facilitation we measured the size of the EMG response 75-125 ms after target appearance relative to the total EMG response in the control (unstimulated) condition (see Methods). We found a significant effect of all stimuli on RVR amplitude (median nerve: F=6.45, P<0.001, Figure 2*B*; vestibular: F=7.07, P<0.001, Figure 3*B*; auditory: F=9.60, P<0.001, Figure 4*B*). There was also a significant effect of target direction on RVR amplitude (median nerve: F=7.29, P=0.002, Figure 2*B*; vestibular: F=9.69, P<0.001, Figure 3*B*; auditory: F=5.67, P=0.005, Figure 4*B*). The general trend of an increase in RVR amplitude was observed with all stimuli and target directions but post-hoc analysis did not identify a specific stimulus latency that was most effective. Similarly, although the majority of subjects showed an increase in RVR with stimuli, this was typically significant in only around half of subjects (median nerve: Figure 2*C*; vestibular: Figure 3*C*; auditory: Figure 4*C*).

**Figure 2.**
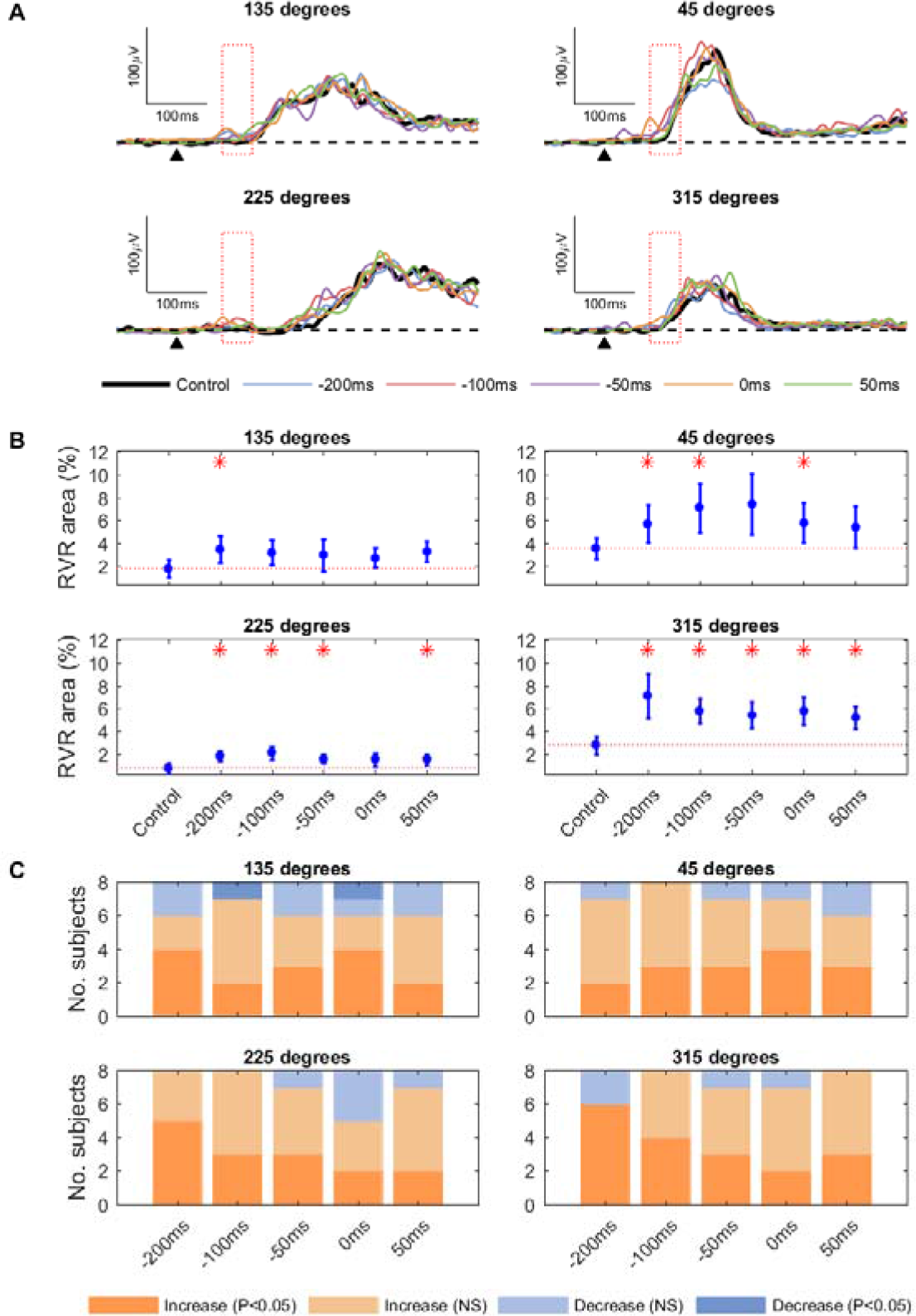
Modulation of RVRs with median nerve stimulation. ***A.*** Mean rectified EMG traces from a single subject showing task-related EMG activity for each median nerve stimulus latency. Each plot represents a different target direction. The black dotted line shows baseline EMG activity. The black arrow indicates target appearance. The red box shows the RVR window (75-125 ms). ***B.*** Mean RVR amplitude (see Methods) averaged across all subjects, displayed for each stimulus condition and target direction. Error bars represent standard error. The red line shows the control condition RVR amplitude, and red asterisks represent a statistically significant (P<0.05) deviation from this. ***C.*** Number of subjects showing an increase or decrease in RVR amplitude with median nerve stimulation, displayed for each median nerve latency and target direction.

**Figure 3.**
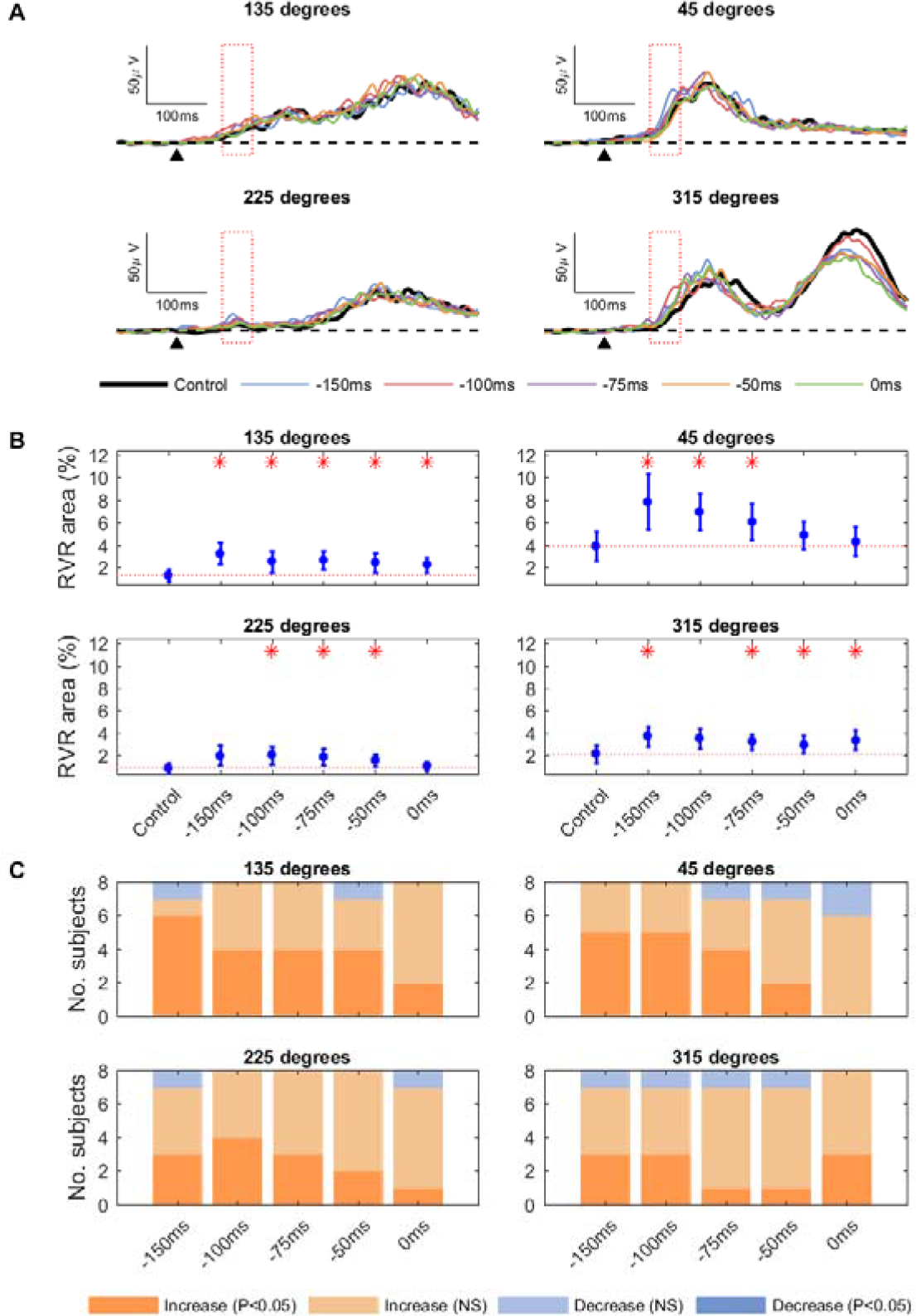
Modulation of RVRs with vestibular stimulation. ***A.*** Mean rectified EMG traces from a single subject showing task-related EMG activity for each vestibular stimulus latency. Each plot represents a different target direction. The black dotted line shows baseline EMG activity. The black arrow indicates target appearance. The red box shows the RVR window (75-125 ms). ***B.*** Mean RVR amplitude (see Methods) averaged across all subjects, displayed for each stimulus condition and target direction. Error bars represent standard error. The red line shows the control condition RVR, and red asterisks represent a statistically significant (P<0.05) deviation from this. ***C.*** Number of subjects showing an increase or decrease in RVR amplitude with vestibular stimulation, displayed for each latency and target direction.

**Figure 4.**
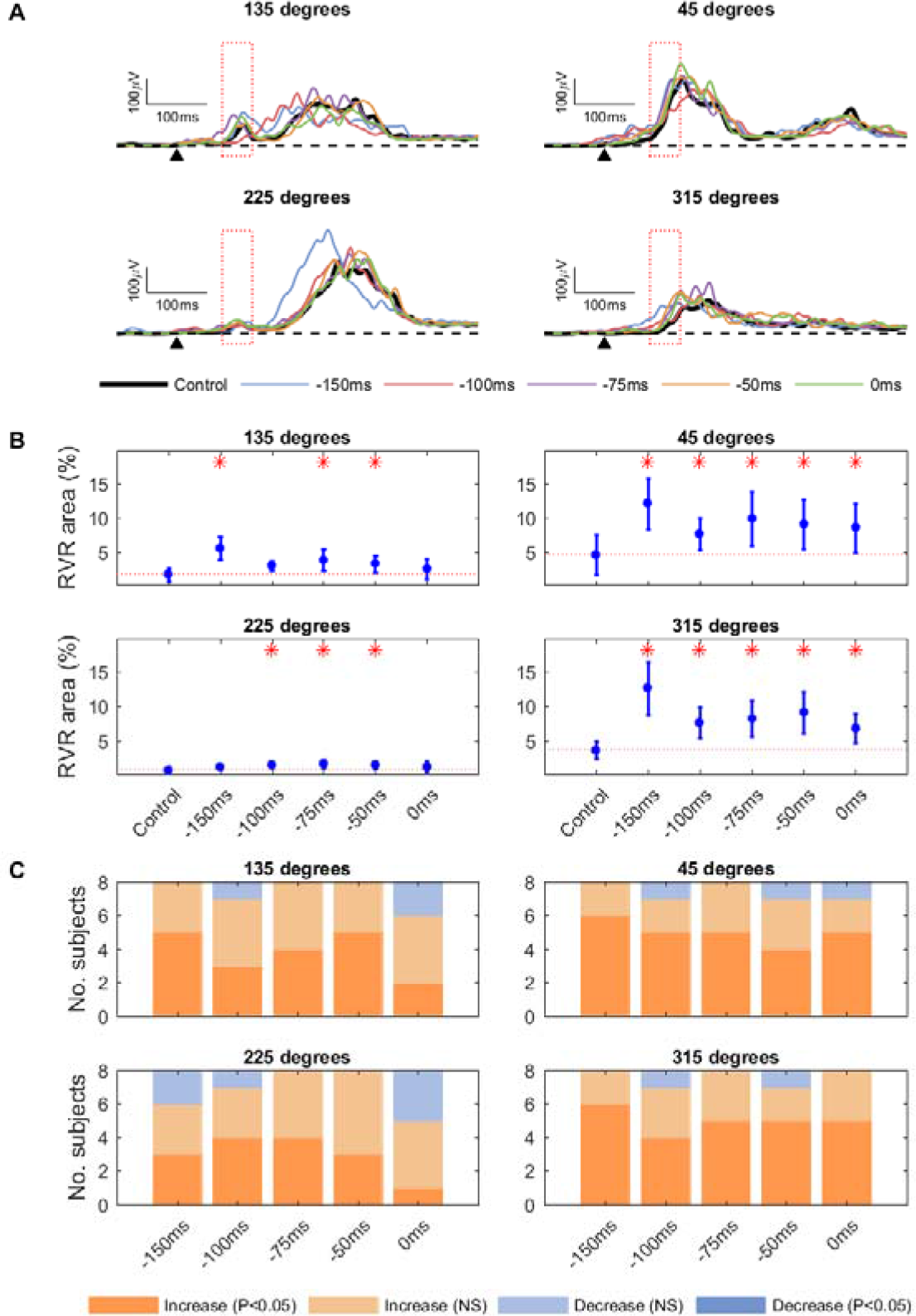
Modulation of RVRs with auditory stimuli. ***A.*** Mean rectified EMG traces from a single subject showing task-related EMG activity for each auditory stimulus latency. Each plot represents a different target direction. The black dotted line shows baseline EMG activity. The black arrow indicates target appearance. The red box shows the RVR window (75-125 ms). ***B.*** Mean RVR amplitude (see Methods) averaged across all subjects, displayed for each stimulus condition and target direction. Error bars represent standard error. The red line shows the control condition RVR, and red asterisks represent a statistically significant (P<0.05) deviation from this. ***C.*** Number of subjects showing an increase or decrease in RVR amplitude with auditory stimuli, displayed for each latency and target direction.

To examine the RVR in isolation from overall changes in EMG activity, we also calculated RVR as a percentage of the total EMG activity of the same single trial, rather than the control condition. This still showed a significant effect on the RVR amplitude of vestibular and auditory stimuli (vestibular: F=4.09, P=0.005; auditory: F=3.25, P=0.016) but not of median nerve stimulation (F=1.48, P=0.222).

### Effects of Stimuli on EMG Latency

EMG onset latency in the control condition was generally in the 75-125 ms range (median nerve: Figure 5; vestibular: Figure 6; auditory: Figure 7), corresponding to the stimulus-locked responses reported by Pruszynski et al. (2010). Pairing target appearance with the different stimuli significantly reduced EMG onset latencies (median nerve: F=6.33, P<0.001, Figure 5*A*; vestibular: F=6.58, P<0.001, Figure 6*A*; auditory: F=12.9, P<0.001, Figure 7*A*). The latency reduction was not uniform across all stimulus timings but instead demonstrated a positive correlation, with the earliest stimulus evoking the shortest latency EMG response (median nerve: Figure 5*B*; vestibular: Figure 6*B*; auditory: Figure 7*B*). Importantly, the reduction in EMG latency did not simply equal the relative stimulus latency. For example, for the 135° target with median nerve stimulation, there was on average a 0.49 ms reduction in EMG latency for every 1 ms that the stimulus timing was advanced (Figure 5*B*). Across all stimuli and target directions, the regression slope was 0.400 ± 0.124 (mean ± SD), which is significantly less than the slope of 1.0 expected if responses simply followed the stimulus timing (P<0.001). Target direction had a significant effect on EMG latency for median nerve (F=3.45, P=0.035) and vestibular stimuli (F=6.21, P=0.004) but not auditory stimuli (F=1.02, P=0.405).

**Figure 5.**
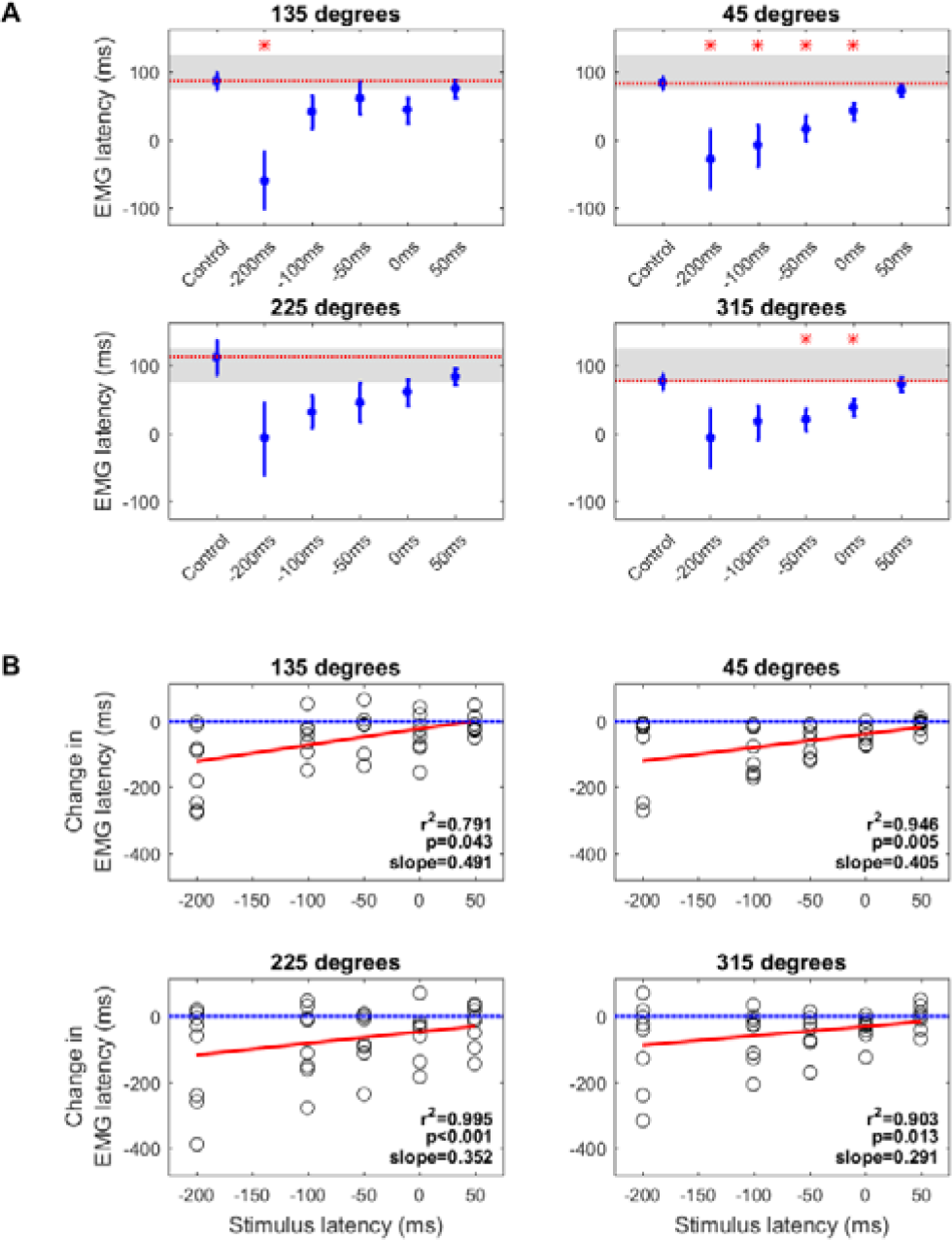
EMG onset latency with median nerve stimulation. ***A.*** Mean EMG latency averaged across all subjects, presented for each target direction (individual plots) and for each median nerve stimulus latency. Error bars represent standard error. The red dotted line shows the EMG latency for the control condition, and the red asterisks represent a statistically significant (P<0.05) deviation from this for each stimulus latency. Grey boxes show the RVR window of 75-125 ms. ***B.*** Correlation of the change in EMG onset latency with stimulus latency. Each point represents the mean change in EMG latency relative to the control condition for one subject in the specified direction. The red line shows the linear regression, with the r^2^ and p values for this displayed on the plot. The blue line represents no change in EMG latency relative to the control condition. Negative values indicate a reduction in EMG latency relative to the control condition.

**Figure 6.**
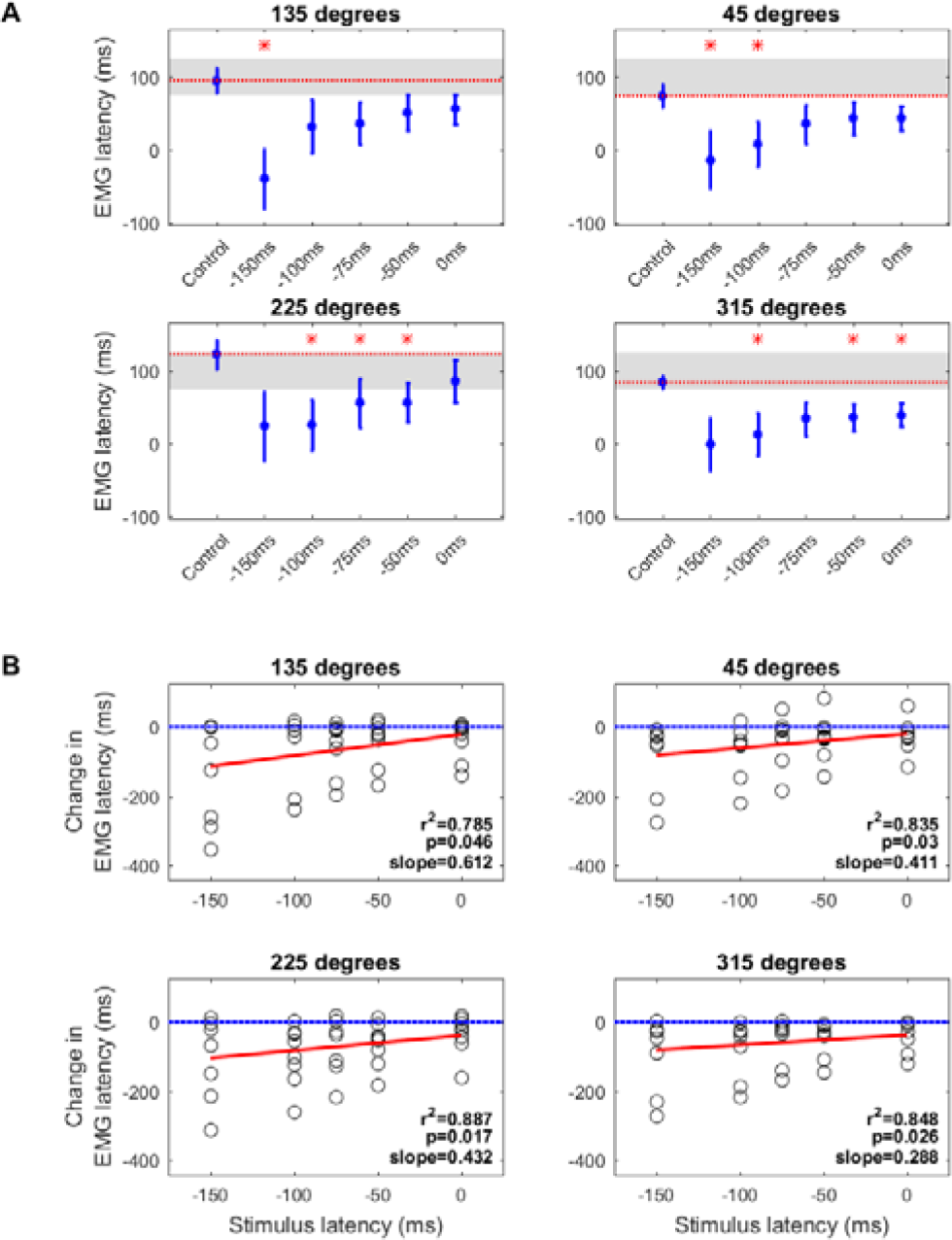
EMG onset latency with vestibular stimulation. ***A.*** Mean EMG latency for averaged across all subjects, presented for each target direction (individual plots) and for each vestibular stimulus latency. Error bars represent standard error. The red dotted line shows the EMG latency for the control condition, and the red asterisks represent a statistically significant (P<0.05) deviation from this for each stimulus latency. Grey boxes show the RVR window of 75-125 ms. ***B.*** Correlation of change in EMG onset latency against stimulus latency. Each point represents the mean change in EMG latency relative to the control condition for one subject in the specified direction. The red line shows the linear regression, with the r^2^ and p values for this displayed on the plot. The blue line represents no change in EMG latency relative to the control condition. Negative values indicate a reduction in EMG latency relative to the control condition.

**Figure 7.**
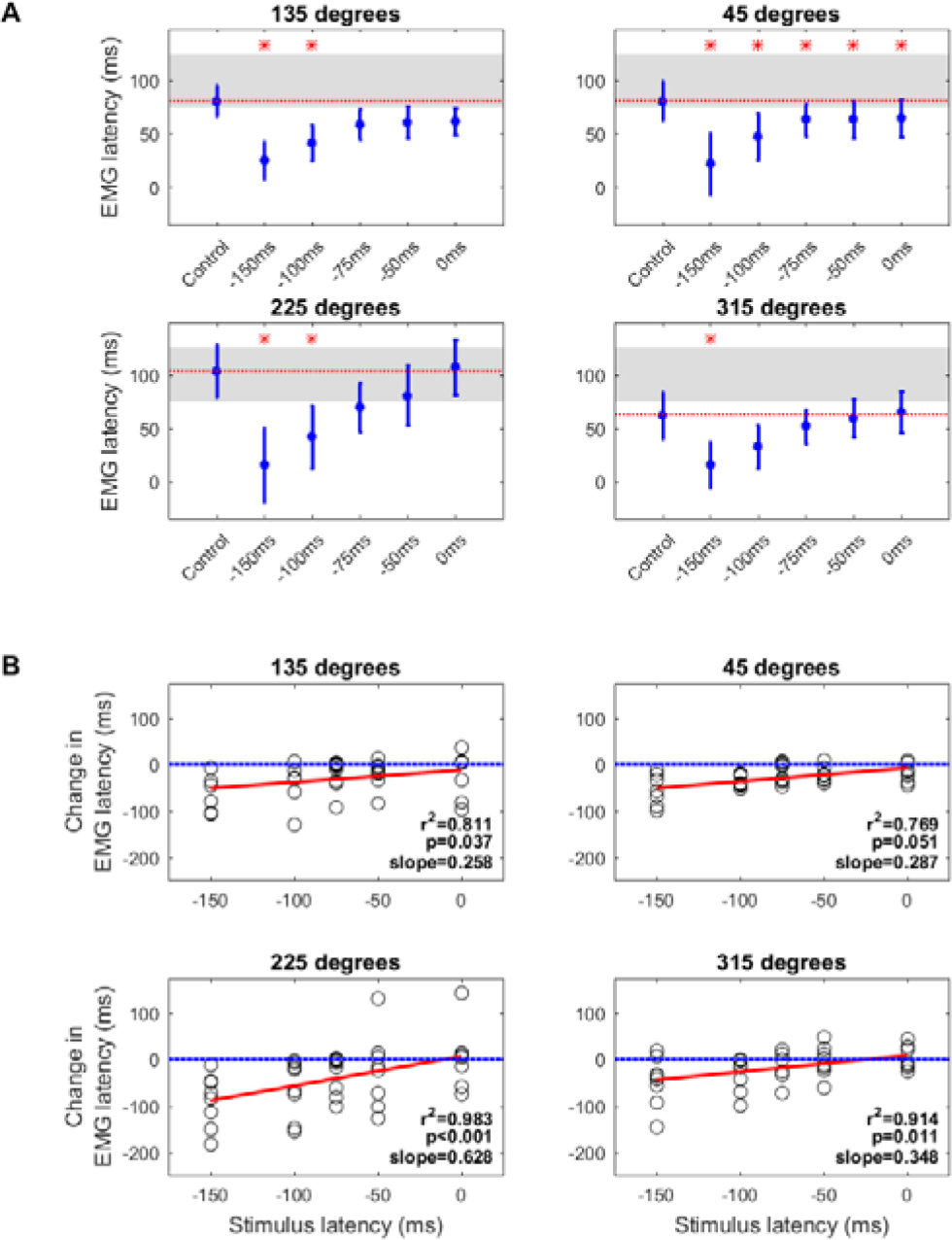
EMG onset latency with auditory stimuli. ***A.*** Mean EMG latency for averaged across all subjects, presented for each target direction (individual plots) and for each auditory stimulus latency. Error bars represent standard error. The red dotted line shows the EMG latency for the control condition, and the red asterisks represent a statistically significant (P<0.05) deviation from this for each stimulus latency. Grey boxes show the RVR window of 75-125 ms. ***B.*** Correlation of change in EMG onset latency against stimulus latency. Each point represents the mean change in EMG latency relative to the control condition for one subject in the specified direction. The red line shows the linear regression, with the r^2^ and p values for this displayed on the plot. The blue line represents no change in EMG latency relative to the control condition. Negative values indicate a reduction in EMG latency relative to the control condition.

### Effects of Stimuli on Total EMG Activity

The effects of the different stimuli were not limited to the early component of the response. There was also a significant effect of all stimuli on the total EMG activity generated in each trial (Figure 8*A*; median nerve: F=4.38; P0.003; vestibular: F=3.31, P=0.015; auditory: F=8.87, P<0.001). This was particularly interesting given that stimulation reduced the time taken to reach target distance (see Task Performance below), thereby shortening the window over which EMG activity was measured. However, it should be noted that the increase in EMG activity showed considerable inter-subject variability (Figure 8*B*).

**Figure 8.**
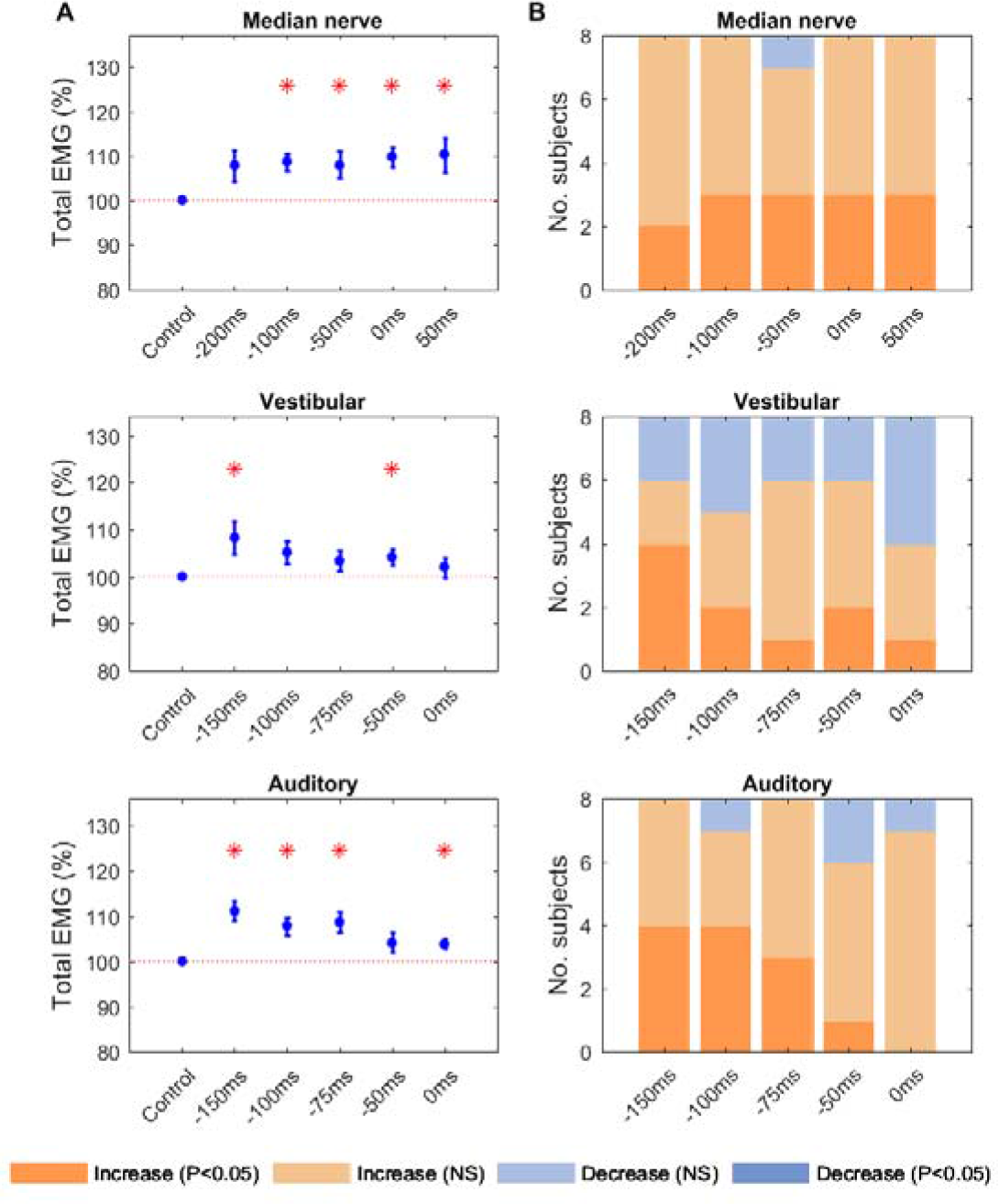
Total EMG activity with median nerve, vestibular and auditory stimuli. ***A.*** Mean total EMG activity for all subjects and target directions, for each stimulus condition. Error bars represent standard error. The red line shows the total EMG activity for the control condition, and red asterisks represent a statistically significant (P<0.05) deviation from this. ***B.*** Number of subjects showing an increase or decrease in total EMG with each stimulus, averaged across target directions and displayed for each stimulus latency.

### Effects of Stimuli on Task Performance

Task performance was assessed by the number of error trials, the time taken to reach the target and the time taken to reach target distance. Although all stimuli had a significant effect on time to reach target distance (red lines, Figure 9; median nerve: F=8.58, P<0.001; vestibular: F=16.0, P<0.001; auditory: F=15.8, P<0.001), indicating improved task performance, this was also associated with a significant increase in error rates (Figure 10; median nerve: F=4.76, P=0.002; vestibular: F=5.88, P<0.001; auditory: F=27.6, P<0.001). Only for median nerve stimulation was there a significant effect on time to reach the target (blue lines, Figure 9; median nerve: F=3.32, P=0.015; vestibular: F=0.29, P=0.916; auditory: F=1.71, P=0.159).

**Figure 9.**
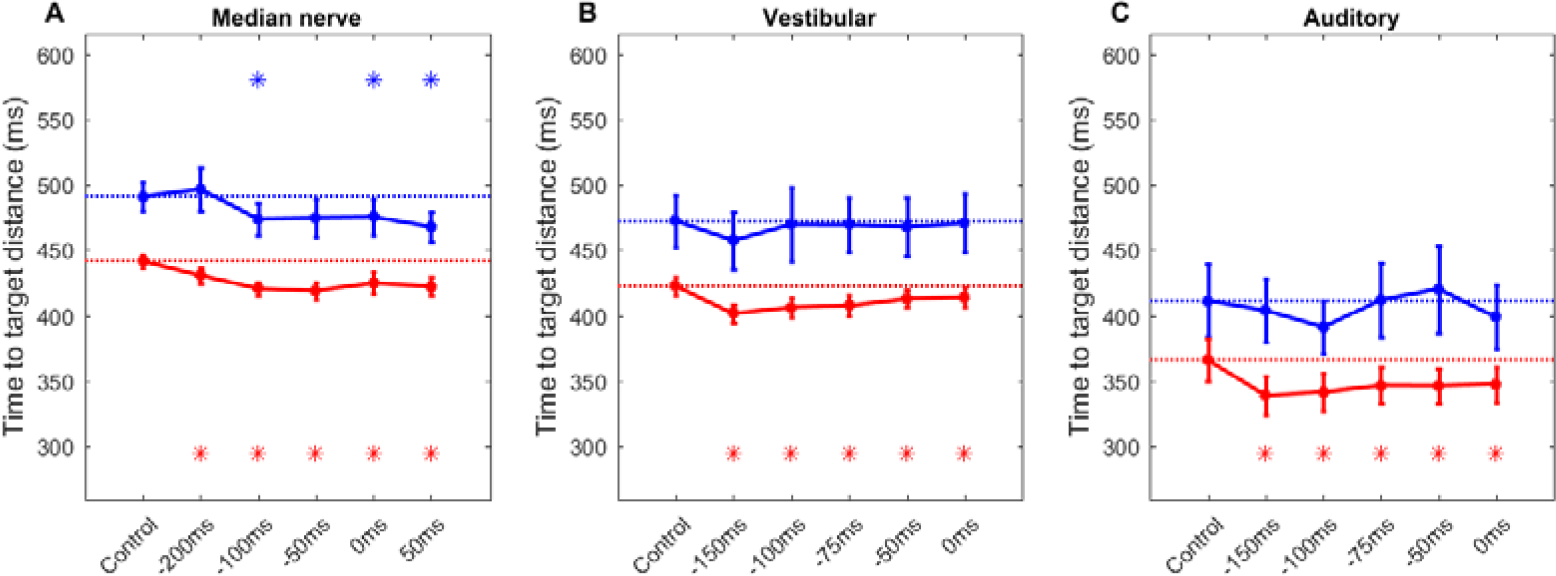
Task performance with median nerve, vestibular and auditory stimuli. Time to reach target (blue) and time to reach target distance (red) averaged across all subjects and target directions, as a function of stimulus timing. ***A***, for median nerve stimulation, ***B***, for vestibular stimulation, ***C***, for loud sound stimulation. Dotted lines represent the control condition and asterisks show a statistically significant (P<0.05) deviation from this. Error bars represent standard error.

**Figure 10.**
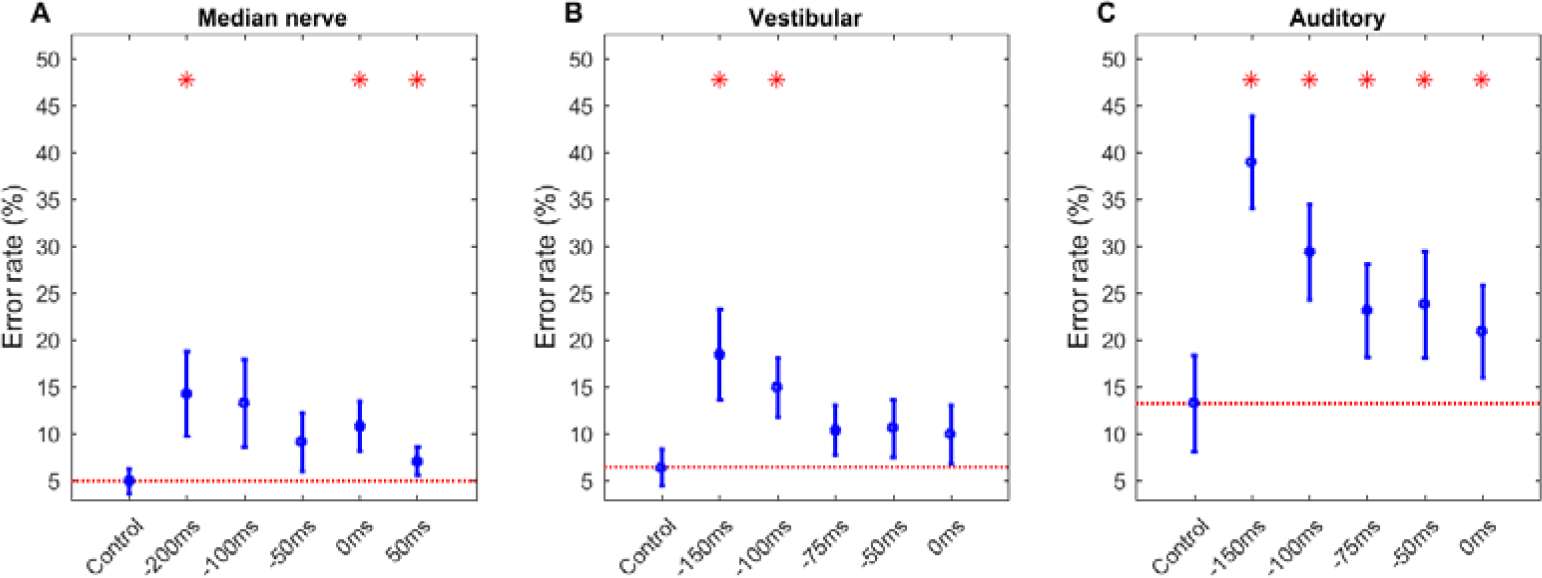
Task error rates. Mean number of errors (trials in which movement was made in the wrong direction) averaged across all subjects and target directions. ***A***, for median nerve stimulation, ***B***, for vestibular stimulation, ***C***, for loud sound stimulation. Dotted lines represent the control condition and asterisks show a statistically significant (P<0.05) deviation from this. Error bars represent standard error.

## Discussion

Increasing evidence suggests that reaching movements are not purely the domain of the cortex but can also be initiated or corrected in a more reflexive manner at short latency. Subcortical structures are an obvious candidate for such visual reflexes (Alstermark et al. 1987; Day and Lyon 2000).

The tecto-reticulospinal tract transforms visual input to motor output via the superior colliculus and RF (Philipp and Hoffmann 2014; Stuphorn et al. 1999; Werner 1993). Since this pathway bypasses the cortex, it is relatively independent of volitional intent (Day and Lyon 2000; Gu et al. 2016) and generates responses at short latencies (Pruszynski et al. 2010). Thus it has been proposed that tecto-reticulospinal output can be recorded by measuring the early component of naturalistic reaching movements made toward visual stimuli. This opens the exciting possibility that reticulospinal excitability can be non-invasively assessed in man.

We paired the reaching task described by Pruszynski et al. (2010) with median nerve, vestibular and auditory stimuli, all of which are known to provide inputs to the brainstem (Irvine and Jackson 1983; Jassik-Gerschenfeld 1966; Ladpli and Brodal 1968; Leiras et al. 2010; Maeda et al. 1979; Mellott et al. 2018; Peterson and Abzug 1975). We found that this resulted in facilitation of the RVR, the short-latency response thought to represent tecto-reticulospinal output, as well as a reduction in EMG onset latency. We propose that both these effects are most likely because the stimuli modulated tecto-reticulospinal excitability.

### Site of Facilitation Effects

The interaction between the various stimuli tested here and the visual input related to target appearance could occur at multiple different levels of the nervous system, but we believe that the cortex is an unlikely site for the RVR facilitation. Although several of the stimuli used can modulate cortical excitability, the effect is largely inhibitory and more dependent on specific timing compared to the facilitation over a wide range of inter-stimulus intervals which we observed. Loud auditory stimuli suppress cortical excitability 30-60 ms after they are delivered (Furubayashi et al. 2000), whilst median nerve stimulation produces both short- (19-21ms; Tokimura et al. 2000) and long-latency inhibition of cortical excitability (200-1000ms; Chen et al. 1999). We are not aware of any reports of the effects of vestibular stimulation on the excitability of upper limb regions of the cortex, although such effects have been reported for the cortical control of neck muscles (Guzman-Lopez et al. 2011). Furthermore, compared to sub-cortical structures, the convergence of sensory inputs onto cortical neurons is less pronounced. Lamarre et al. (1983) reported that although 30% of M1 cells recorded responded to light, sound or torque pulses, only 10% responded to multiple stimuli and no summation was apparent when these stimuli were combined.

Assuming that the RVR is carried over a tecto-reticulospinal route, stimulus interactions could occur at each stage of this pathway. The superior colliculus receives a wide range of inputs, including from the limbs (Jassik-Gerschenfeld 1966), vestibular system (Maeda et al. 1979) and auditory system (Mellott et al. 2018). Convergence and facilitation of the RVR is thus possible even at this early stage of processing. In addition, numerous studies show multimodal responses in the RF, to inputs including auditory, visual, somatosensory and vestibular stimuli (Martin et al. 2010; Miller et al. 2017; Oliveras et al. 1990; Oliveras et al. 1989; Wepsic 1966). The functional relevance of this sensory convergence is apparent in the startle reflex, which is mediated via the RF (Brown 1995) and is more effectively elicited by multimodal summation of tactile, auditory and vestibular inputs than intramodal temporal summation (Yeomans et al. 2002). Furthermore, paired delivery of auditory clicks and peripheral electrical stimulation can generate lasting changes in the long-latency stretch reflex (Foysal et al. 2016), which may partially depend on reticulospinal outputs (Soteropoulos et al. 2012). Further support for a role of the RF comes from the wide range (250 ms) of stimulus timing which was capable of facilitating the RVR. This suggests that stimuli had a rapidly-induced but long-lasting effect on excitability. It is known that appropriate stimulation can increase the firing rate of cells in the nucleus reticularis gigantocellularis for extended periods (Martin et al. 2010). Even in anaesthetized macaques, auditory stimuli can increase RF firing rates for up to 25 ms (Fisher et al. 2012). Combined, these studies provide strong support for the brainstem as a site of multisensory integration and thus a likely locus for the facilitation of RVRs.

It is also possible that the RVRs were facilitated by the different stimuli at the level of the spinal cord. Many spinal interneuron systems show extensive convergence of descending inputs from vestibulospinal, reticulospinal and corticospinal tracts (Illert et al. 1981; Illert et al. 1977; Krutki et al. 2017; Riddle and Baker 2010; Suzuki et al. 2017) as well as from peripheral afferents (Pierrot-Deseilligny and Burke 2012). Loud sounds may excite the vestibular apparatus (Watson and Colebatch 1998) as well as the reticular formation, hence both the auditory and vestibular stimuli could be interacting with descending reticulospinal commands within the spinal cord (Yeomans et al. 2002). However, spinal interactions between converging stimuli tend to be highly specific for timing (Pierrot-Deseilligny and Burke 2012). Furthermore, at least for the well-characterized C3-C4 propriospinal system, facilitation is typically followed by feedback suppression, which makes the demonstration of interactions highly dependent on selection of an appropriate stimulus intensity. Whilst weak stimuli show no effect, strong stimuli above motor threshold may generate overlapping suppression and facilitation and also fail to generate consistent changes in the test response (Malmgren and Pierrot-Deseilligny 1987; Mazevet and Pierrot-Deseilligny 1994). By contrast, we found robust effects using relatively strong median nerve stimuli (intensity twice motor threshold) at a wide range of stimulus timings. Although we cannot rule out some contribution of convergence at spinal interneurons for RVR facilitation in our results, this is likely to be less important than convergence within the brainstem.

Finally, we must consider whether the effects which we observed were generated by changes at the level of the motoneuron. It is known that motoneuron excitability increases for several hundred milliseconds after a warning cue (Komiyama and Tanaka 1990; Rossignol and Jones 1976). Such an effect could explain the increase in total EMG produced during the task (Figure 8). However, we found that there was a disproportionate facilitation in RVR, which increased when expressed as a fraction of the total EMG. This implies a mechanism which is selective for this part of the response, rather than merely raising all muscle activity. Changes in motoneuron excitability alone cannot therefore explain our findings.

### Latency Effects

In addition to the facilitation of RVRs, EMG onset latency was reduced by all stimuli which we tested. This is reminiscent of a StartReact phenomenon whereby startling stimuli reduce reaction time by early release of a prepared motor program (Valls-Sole et al. 1999). StartReact requires that the movement is known in advance such that it can be prepared and stored; StartReact effects are absent in choice reaction tasks (Carlsen et al. 2004b). Furthermore, the response profile with StartReact should be unaltered (Carlsen et al. 2004a; Dean and Baker 2017; Valls-Sole et al. 1999). Given that we used a choice reaction task, showed an increase in total EMG activity, and observed the latency shift with all three stimuli tested (and not just the loud sound), we cannot simply characterize the phenomenon which we describe as a StartReact effect.

An alternative hypothesis for the reduction in onset latency is intersensory facilitation (Hershenson 1962), which is the speeding up of motor preparation by accessory stimuli (Nickerson 1973; Schmidt et al. 1984). Although intersensory facilitation is observed in choice reaction tasks (Schmidt et al. 1984) and thus provides a more appropriate model for our data, it is difficult to reconcile our observation that EMG activity can increase prior to the visual target appearance with the model of hastened motor preparation. Furthermore, previous reports have shown accessory stimuli to produce the shortest latency responses when delivered with or following the imperative stimulus (Maslovat et al. 2015; Nickerson 1970; Terao et al. 1997), whereas we found the earliest stimulation most effective in reducing EMG onset latency. Together, these findings suggest that the latency reduction seen here was not generated by the cortically-mediated intersensory facilitation previously described in the literature. This likely reflects the lack of cortical involvement in the RVR.

Reynolds and Day (2007) interacted a visually-cued task with auditory stimulation, and observed a response latency shift. They suggested that this interaction occurred at the caudal pontine RF, leading to faster visuomotor processing. Although a similar mechanism may be at play in our task, it cannot account for increases in muscle activity before the imperative stimulus. This apparently premature increase in muscle activity may instead result from the long-lasting non-specific increase in motoneuron excitability generated by warning cues (Komiyama and Tanaka 1990; Rossignol and Jones 1976). We observed the highest error rates with the earliest stimuli, indicating that the heightened state of readiness increased the likelihood of subjects responding prematurely, before they had determined the correct movement direction. This is likely to be a different effect from the enhancement of the true RVR, which starts around 75 ms after target onset.

### Conclusion

In conclusion, we used a choice reaction reaching task to show that stimuli delivered across a range of latencies can significantly reduce reaction times in a proximal muscle and facilitate short-latency responses. We propose that this reflects modulation of tecto-reticulospinal excitability. Given the wealth of sensory information received by the superior colliculus and RF, it is possible that these structures act as a site of multisensory integration; appropriate pairing of inputs may provide a means of modulating their output. In the context of accumulating evidence supporting a role of the RST in functional recovery, and the limitations of recovery, after corticospinal lesions (Baker 2011; Baker et al. 2015; Dewald et al. 1995; McPherson et al. 2018; Zaaimi et al. 2012; Zaaimi et al. 2018), we tentatively suggest that the ability to influence reticulospinal excitability non-invasively with such techniques may find clinical utility.

